# Sex and the facilitation of cued fear by prior contextual fear conditioning in rats

**DOI:** 10.1101/2024.05.23.595599

**Authors:** Katherine Vazquez, Kehinde E. Cole, Ryan G. Parsons

**Author notes:** Corresponding Author: Ryan G. Parsons, Department of Psychology, 100 Nicolls Rd., Stony Brook, NY, 11794.

## Abstract

Previous studies have shown that the formation of new memories can be influenced by prior experience. This includes work using pavlovian fear conditioning in rodents that have shown that an initial fear conditioning experience can become associated with and facilitate the acquisition of new fear memories, especially when they occur close together in time. However, most of the prior studies used only males as subjects resulting in questions about the generalizability of the findings from this work. Here we tested whether prior contextual fear conditioning would facilitate later learning of cued fear conditioning in both male and female rats, and if there were differences based on the interval between the two conditioning episodes. Our results showed that levels of cued fear were not influenced by prior contextual fear conditioning or by the interval between training, however, females showed lower levels of cued fear. Freezing behavior in the initial training context differed by sex, with females showing lower levels of contextual fear, and by the type of initial training, with rats given delayed shock showing higher levels of fear than rats given immediate shock during contextual fear conditioning. These results indicate that contextual fear conditioning does not prime subsequent cued fear conditioning and that female rats express lower levels of cued and contextual fear conditioning than males.

---

Several recent studies have shown that prior fear conditioning can enhance the acquisition of subsequent fear learning (Parsons and Davis, 2012; Rashid et al., 2016; Parsons, Walker, and Davis, 2016; Lee et al., 2018; Parsons, 2018; Cole and Parsons, 2023). These facilitation-like effects are time dependent such that the enhancement of subsequent learning is optimal when the two training experiences are separated by several hours, and weaker when they are separated by shorter (i.e. 4 minutes) or longer (i.e. 7 days) time intervals (Parsons & Davis, 2012; Lee et al., 2018; Cole and Parsons, 2023). This work is conceptually similar to other behavioral and cellular studies on synaptic tagging, metaplasticity, and memory allocation, suggesting that the ability of learning to alter subsequent memory formation is a prevalent feature of information storage in the brain (Sehgal, et al., 2013; Josselyn and Frankland, 2018; Nomoto and Inokuchi, 2018). Although these facilitation effects are evident in cued fear conditioning in male rodents, little is known about whether females exhibit similar facilitation effects and whether contextual fear conditioning facilitates subsequent cued fear learning in a similar manner.

Determining whether there are sex differences in the ability of initial learning to facilitate subsequent fear conditioning is important for understanding basic learning and memory processes and it might also have relevance to post-traumatic stress disorder (PTSD). This is because fear conditioning likely underlies some aspects of the development of PTSD following trauma (Glover et al., 2012; Inslicht et al., 2013; Parsons and Ressler, 2013), and women are about twice as likely to develop PTSD as compared to men (Kilpatrick et al., 2013). Moreover, there is evidence that the incidence and severity of PTSD are on average proportional to the number of traumatic events (Breslau et al., 1999; Brewin et al., 2000; Suliman et al., 2009).

Thus, experimental paradigms that seek to understand how prior fear conditioning experiences can influence subsequent learning might be relevant to the mechanisms supporting cumulative trauma.

The goal of the present study was to determine whether the effect of prior learning on subsequent fear conditioning was similar in females and males and to determine if contextual fear conditioning is able to prime the acquisition of subsequent cued fear. To test this, the present study assessed male and female rats exposed first to a contextual fear conditioning procedure in which rats of both sexes received two shocks either immediately after placement in the conditioning chamber or after a delay of several minutes, and then at variable intervals underwent a second conditioning procedure in which there was a single pairing of light and shock. We used the immediate versus delayed shock procedure because it supports contextual fear conditioning in only one of the groups (i.e. delayed shock) (e.g. Fanselow, 1990; Rudy and O’Reilly, 2001) while equating exposure to the aversive and contextual stimuli. This is important because it allows us to determine if it is associative learning specifically that supports the facilitation of subsequent fear memory formation or if non-associative processes (i.e. sensitization) are responsible. After exposure to the two conditioning procedures, fear of the light cue and context in which initial training occurred was tested in separate sessions.

## Method

### Subjects

144 adult Sprague Dawley rats (72 females and 72 males) served as subjects. Rats were ordered from Charles River Laboratories and delivered at approximately 8 weeks of age. Rats were given a week to acclimate to the facilities prior to testing. Rats were given free access to food and water and were housed in pairs on a 12-hour light/dark cycle, with lights on at 7 am and lights off at 7 pm. Testing occurred during the light portion of the cycle. All experimental procedures involving the rats were approved by the Stony Brook University Institutional Animal Care and Use Committee and followed the guidelines for the care and use of laboratory animals set by the National Institutes of Health.

### Behavioral Apparatus

#### Startle Chambers

Baseline acoustic startle, visual fear conditioning, and visual fear testing occurred in startle chambers (Startle Monitor II, Kinder Scientific, Poway, CA, USA). Baseline startle occurred with rats placed in the startle chambers inside of restraint cages with plastic floors and a stainless-steel rod cover. During visual fear conditioning and visual fear testing, rats were placed in the startle chambers inside of restraint cages with stainless-steel shock grid floors and walls made of Plexiglass. Startle responses from the rats were provoked by playing a 50ms, 95 dB white noise burst through speakers placed on the startle chamber ceilings. Startle amplitude was measured in Newtons (N), and startle amplitude was recorded as the maximum N during a 500msec window following the startle stimulus. Shocks (0.4mA/0.5sec) were delivered through the shock grid floors of the restraint cages. For visual fear conditioning and visual fear testing, rats were shown a light cue (82 lux/4.0 sec) delivered through an LED panel mounted on the ceiling of the chambers. Cages were scented with 70% EtOH during testing.

#### Freezing Chambers

Context conditioning and context testing occurred in freezing chambers (Clever Systems Inc., Reston, VA, USA) that were housed in sound attenuating boxes. Chambers consisted of Plexiglass walls and shock grid floors and were wiped down with 10% acetic acid during both the training and testing sessions. 28-V, incandescent, house light bulbs illuminated the inside of each chamber. Cameras positioned overhead recorded the animals’ behavior during training and testing sessions. A computer program (FreezeScan 2.0) was used to control the delivery of the stimuli and to score animals’ freezing behavior. Shocks of 1.0mA were delivered through the shock grid floors of the freezing chambers.

### Behavioral Procedures

Each animal was handled for 5 minutes per day for 7 days before the experiment began. The first 4 days of handling took place in the colony room while rats were carted into the testing room and handled on the final 3 days. The first two days of each experiment were dedicated to baseline startle. Baseline startle was assessed by placing the rats into the startle chambers in restraint cages and subjecting them to a five-minute period in which no startle stimuli were presented. Following this acclimation period, the rats were subjected to 30 white noise bursts at 95db, with each burst lasting for 50 milliseconds (30s ITI). The startle amplitude for each rat was averaged across each day and the average startle amplitude was used to match rats into groups so that within-sex the average startle amplitude was roughly equal across groups.

The following day, rats underwent contextual fear conditioning in the freezing chambers. Rats were matched into immediate and delayed groups. Rats in the immediate shock group were placed in the freezing chambers and exposed to two 1mA foot shocks (2s ITI) within 10 seconds of being placed inside of the chambers and were returned to their home cages after 3 minutes in the chamber. Rats in the delayed shock group were placed in the freezing chambers and were given two 1mA foot shocks (3s ITI) after being in the chambers for 6 minutes. Rats were returned to their home cages after 11 minutes inside the conditioning chamber. The conditioning protocol used 2 shocks rather than a single shock as pilot experiments from our lab showed negligible levels of freezing during testing in rats that had received a single delayed shock during training.

One hour, one day, or one week after context fear conditioning, rats underwent visual fear conditioning in the startle chambers. All rats were placed in restraint cages inside of the startle chambers and after five minutes were exposed to 4.0 seconds of light followed by a 0.4mA foot shock. One minute after the shock, the rats were returned to their home cages. Forty-eight hours after visual fear conditioning, rats underwent visual fear testing. During this test session, rats were placed in the startle cages and, after a five-minute acclimation period, were exposed to 30 white noise bursts at 95db, with each burst lasting for 50 milliseconds (30s ITI). Following this, rats were subjected to ten trials of 4.0 seconds of light that co-terminated with a 95db white noise burst. Each light-startle trial was followed by three 95db white noise bursts without light before the next light trial began. Rats were returned to their home cages following the final trial. On the final day, rats underwent context testing in the freezing chambers. Rats were placed in the chambers for 10 minutes without being subjected to any startle stimuli.

### Data Analysis

Statistical analyses were performed using GraphPad Prism software. All rats underwent 2 days of baseline startle, and baseline startle averages were obtained by averaging each rat’s startle amplitude across both days. These values were used to match rats into groups with equivalent mean startle amplitude within each sex. Shock reactivity during the visual fear conditioning session was measured as the peak change in force that occurred during the shock period. Fear-potentiated startle was determined by obtaining a difference score by subtracting the startle alone trial averages from the average startle response during the light CS and expressing this value as a percent change score. Fear during the intertrial interval periods during the light test session (i.e. ‘ITI fear’) was determined by subtracting individual rats’ average acoustic startle during the 30 pre-CS startle trials from the 30 startle trials in between light-startle trials. Freezing during contextual fear conditioning was calculated by taking an average of the percent time spent freezing after delivery of the final shock. During the context test, time spent freezing was averaged for each rat over the 10 minutes of testing. Three-way ANOVAs with sex, training type, and time interval as factors were used to test for differences. When analyzing each sex separately, two-way ANOVA was used with group and time interval as factors. Sidak’s post hoc tests were used to assess the significance of differences between groups. Finally, Pearson’s r was used to correlate individual rats’ fear-potentiated startle during testing with freezing behavior during the context testing session. Results were considered significant if *p* < 0.05.

## Results

First, a three-way ANOVA was performed on baseline startle averages using training type, sex, and interval as the factors. The ANOVA revealed that there was a significant difference in baseline startle averages between males and females, F (1, 132) = 6.79, *p* < .05, with males showing higher average amplitude startle (**Figure 1B**). There were no other significant differences between groups and no interactions. This finding is consistent with our prior observations (Russo and Parsons, 2021; Cole and Parsons, 2023) and may be related to differences in weight between males and females. We also used a three-way ANOVA to compare freezing levels after the final shock in rats given contextual fear conditioning. Results showed that there was a significant effect of training type F (1, 132) = 82.00, *p* < .0001, and a significant sex effect F (1, 132) = 9.14, *p* < .01 with delayed shock rats showing higher levels of freezing than immediate shock rats and males showing higher levels of freezing than females (**Figure 1C**). Finally, we compared shock reactivity during the visual fear conditioning session using 3-way ANOVA. Results from this analysis (**Figure 1D**) showed no main effects and no interactions between groups, indicating that the differences in contextual fear are not the product of differences in processing of the shock.

**Figure 1.**
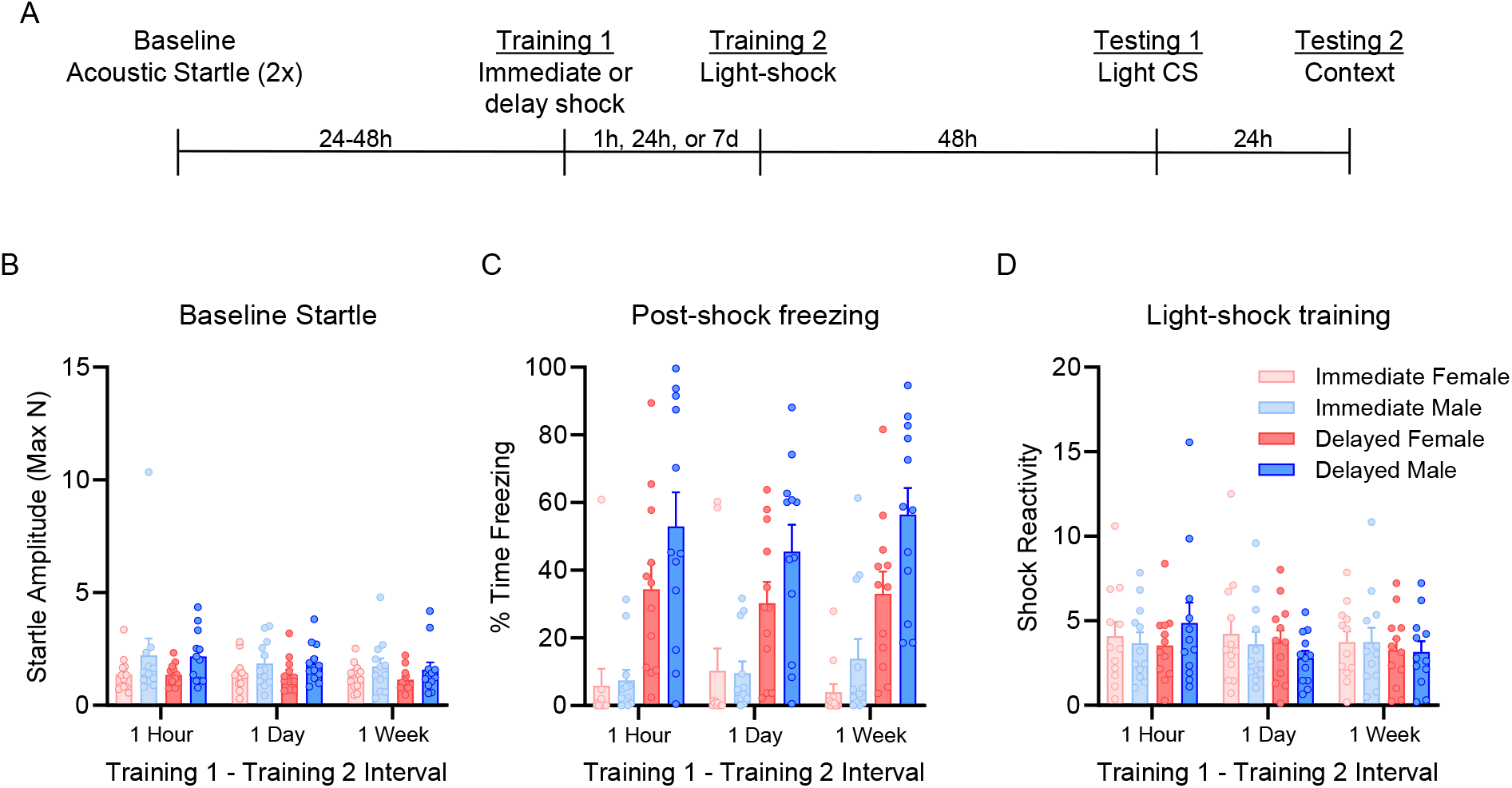
(A) Timeline of behavioral testing. After two days of baseline startle testing, female (N = 72) and male (N =72, N=12/group) rats were given an initial contextual fear conditioning session (Training 1) consisting of 2 shocks presented either within seconds after placement into the conditioning chamber (immediate) or after a 6-minute acclimation period (delayed). 1 hour, 1 day, or 1-week later rats received a second conditioning experience consisting of a single light-shock. Long-term memory to the context and light cue was tested 48 h later, with the light cue tested first and the context the test the next day. (B) Males showed higher average baseline startle than female rats (*p* < .05). (C) Freezing behavior in the 2-minute period after shock in which there was a significant effect of sex (*p* < .01) and training type (*p* < .0001). (D) No significant differences were observed in shock reactivity during visual fear conditioning. Error bars = +/-SEM; Dots = individual means.

Next, levels of fear-potentiated startle during the light CS test were compared using a three-way ANOVA using training type, sex, and interval as the factors. Results showed that there was a main effect of sex F (1, 132) = 5.412, *p* < .05, with males showing higher levels of fear-potentiated startle (**Figure 2A**). There were no other significant main effects and no interaction between factors. Next, we used 3-way ANOVA to analyze fear during the intertrial interval during the light testing session. Results from this test showed a significant effect of sex F (1, 132) = 5.618, *p* < .05, with males showing higher levels of fear during ITI than females (**Figure 2B**). Again, there were no other main effects and no interactions. Finally, we computed fear during the context test session by taking the average time each rat spent freezing during the test session and subjecting these data to a three-way ANOVA with training type, sex, and interval as factors. Results showed a significant difference between males and females, F (1, 132) = 8.579, *p* < .01, with males showing higher freezing levels than females (**Figure 2C**). This analysis also revealed a significant effect of training type, with the delayed group spending more time freezing during the context test than the immediate group F (1, 132) = 38.26, p < .0001. There were no interactions among factors.

**Figure 2.**
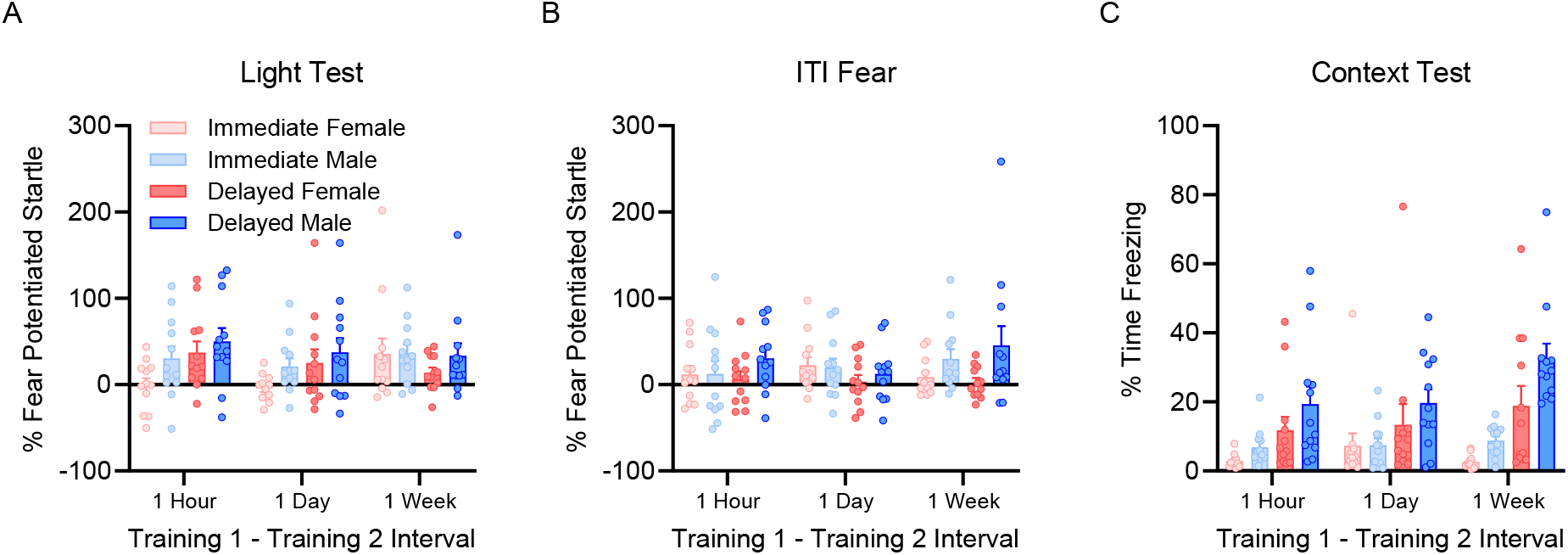
Fear to the light CS, the intertrial interval period of the light test, and to the context. Male rats showed significantly higher (*p* < .05) fear-potentiated startle to the light cue during the light CS test compared to females (A). Fear during the intertrial interval was also higher in males than in females (*p* < .05) during the light CS session (B). Freezing during the context test session was significantly higher in males than in females (*p* < .01) and in rats trained with delayed shocks as compared to rats given immediate shock training (*p* < .0001) (C). Error bars = +/-SEM; Dots = individual means.

To uncover any patterns not evident when the data were combined, we disaggregated the data by sex and performed 2-way ANOVA on the results from both testing sessions. During the light CS test, there were no main effects in females, however, there was an interaction between interval and training type, F (2, 44) = 3.718, *p* < .05 (**Figure 3A**). Sidak’s post hoc tests showed that there was a significant difference (p < .05) between immediate and delayed shock females trained with 1 hour between contextual and visual fear conditioning. No other comparisons were significant in females. In males, there were no main effects or interactions between factors during the light CS test (**Figure 3D**). During the ITI of the light test session, results from a 2-way ANOVA showed no main effects or interactions in either sex (**Figures, 3B, E**). Finally, a 2-way ANOVA on data from the context test session showed a significant effect of training type in females F (1, 22) = 10.15, *p* < .01 (Figure 3C), as well as in males, F (1, 22) = 59.88, *p* < .0001 (**Figure 3F**). There was no interaction between training type and interval in either females or males on data from the context test.

**Figure 3.**
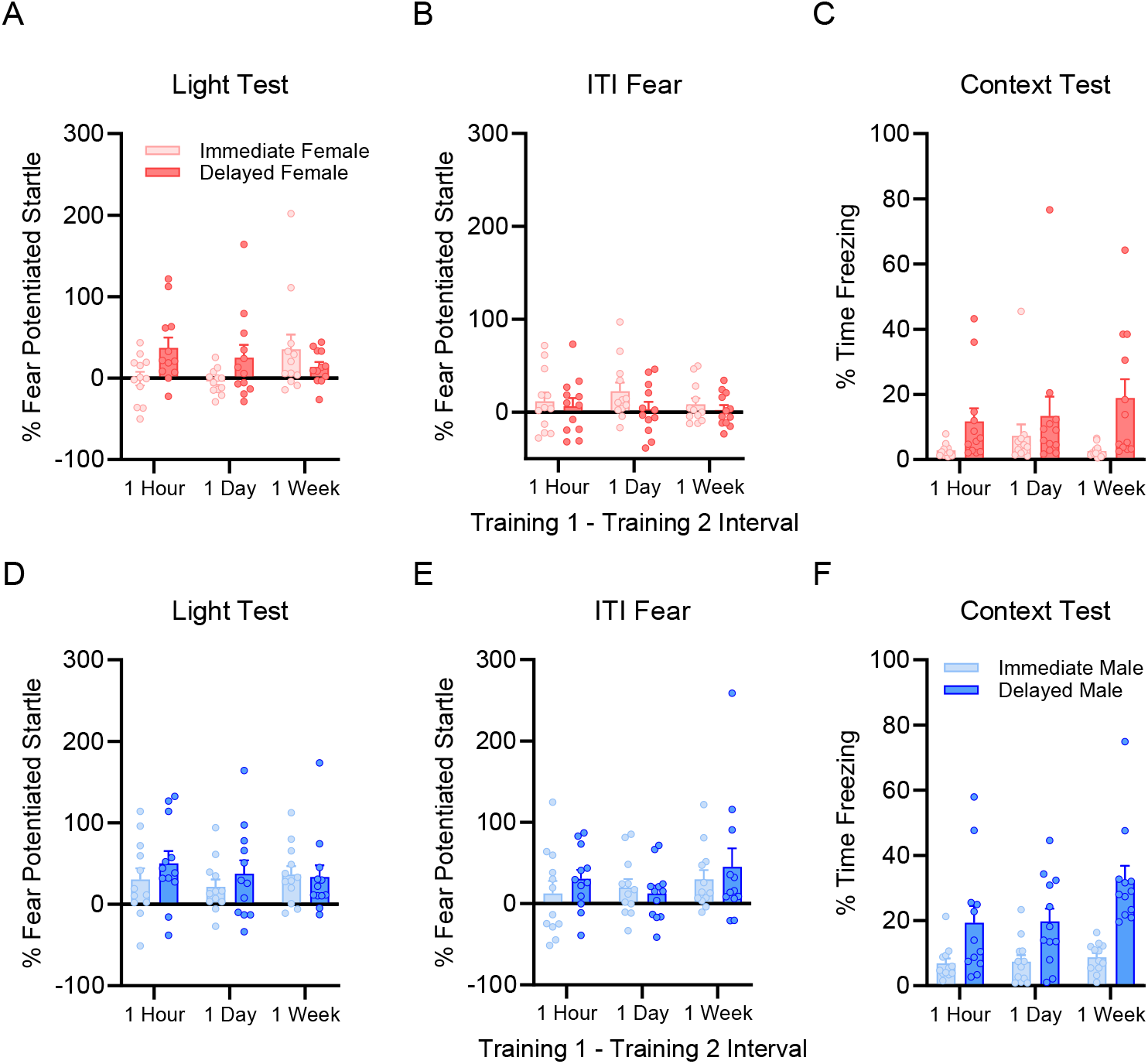
Fear to the light CS (A, D), during the ITI (B, E), and in the conditioning context (C, F) in females (A-C) and males (D-F). During the light CS test in females (A), there was no effect of interval or training type, but there was interaction between the two factors (*p* < .05). There were no significant differences in males, and there were no significant differences in either sex in fear during the ITI (B, E). During the context test, delayed shock animals showed higher levels of fear in both sexes (C, F), but there was no effect of training interval and no interaction.

Finally, in our prior work we observed that for some conditions, high fear of the initially trained cue was associated with lower fear of the subsequently trained cue in individual rats (Lee et al., 2018). To determine if there was a relationship between levels of fear to the light cue and fear to context in this data set, we used Pearson’s r to test for a correlation between individual rats’ levels of fear to the two cues within each condition. No significant correlations were observed in female rats in any of the conditions (**Figure 4A-F**), or in male rats that had received immediate shock (**Figure 4G-I**). However, in males that received delayed shock followed 1 day later by visual fear conditioning there was a significant negative correlation between freezing levels during the context test and fear-potentiated startle to the light (**Figure 4K**). Fear to the light cue did not correlate with context fear in males trained with delayed shock at the 1-hour or 7-day intervals (**Figure 4J, L**).

**Figure 4.**
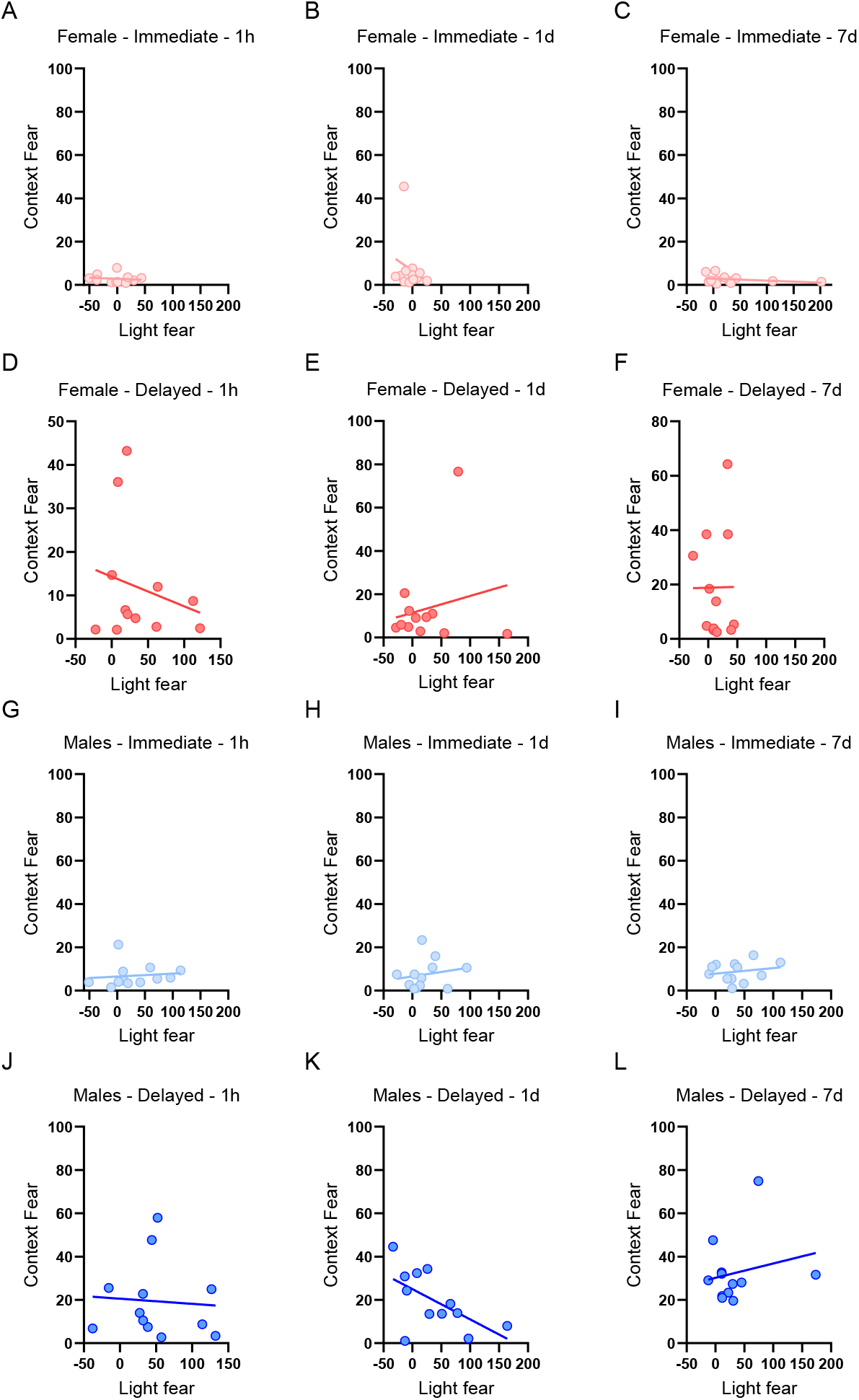
The relationship between levels of fear to the context in which initial training occurred and fear to the light CS for each subject was separated by group (A-K). There was a significant negative correlation between levels of fear to the light CS and fear to the context in males that were trained with the two conditioning events separated by 1 day. No other significant correlations were observed.

## Discussion

This study sought to determine if contextual fear conditioning would facilitate subsequent visual fear conditioning and whether the performance of males and females would differ. Rats were trained initially with either delayed or immediate presentations of shock upon exposure to a conditioning context and then at varying intervals later were exposed to a single pairing of a light cue with shock. Memory for the context and light cue were then tested separately. Consistent with several prior studies (Fanselow, 1990, Rudy and O’Reilly, 2001; Wiltgen et al., 2001), delayed shock rats exhibited higher levels of contextual fear than rats exposed to immediate shocks during training. Yet, despite this difference in the initial learning of contextual fear, there were no differences between the immediate and delay shock groups in subsequent learning of a visual fear conditioning task. These findings suggest that contextual fear conditioning does not prime subsequent learning of visual fear conditioning, which is consistent with recent work from our lab (Cole and Parsons, 2023).

Our conclusion that contextual fear conditioning did not prime subsequent learning is based on the finding that delayed shock rats’ fear to the visual cue did not differ from those given immediate shock during context conditioning, even though there was a robust difference in the levels of contextual fear between these two groups. One alternative explanation is that the lack of a difference in the facilitation of cued fear between the immediate and delay groups is that exposure to the shock during contextual fear conditioning resulted in a non-specific enhancement of fear to the light CS (i.e. pseudoconditioning or sensitization) in both groups. Although we did not have a control to distinguish the facilitation of subsequent learning from pseudoconditioning in the present study, our prior work has tested this directly and found that responding to the light CS is not simply the result of sensitization or pseudoconditioning produced by exposure to the shock during initial learning (Parsons and Davis, 2012; Cole and Parsons, 2023). In addition, the overall level of fear-potentiated startle to the light cue in the present study is relatively low compared to our prior work (Lee et al., 2018; Cole and Parsons, 2023) where auditory fear conditioning preceded visual fear conditioning, providing further support to the conclusion that contextual fear conditioning has little effect on the acquisition of subsequent visual fear conditioning.

Fear-potentiated startle to the light cue did not differ based on whether the rats experienced immediate or delayed training prior to visual fear conditioning, however, there was a significant sex difference with females showing lower fear-potentiated startle to the light cue. In our prior work (Cole and Parsons, 2023) we found lower levels of fear to the light in females when visual fear conditioning was preceded by auditory fear conditioning, although this effect was specific to rats that had been trained with a 24-hour interval. Results from prior studies using more traditional designs in which rats of both sexes are trained with a single conditioning experience are mixed, with some work showing lower fear-potentiated startle in females, and others showing higher fear-potentiated startle in females or no differences (de Jongh et al., 2005; Voulo and Parsons, 2017; Zhao et al., 2018; Olivera-Pasilio and Dabrowska, 2023). Our results also indicate that males showed higher levels of contextual fear than females, which is consistent with a number of prior studies (Maren et al., 1994; Wiltgen et al., 2001; Chang et al., 2009; Gresack et al., 2009; Barker & Galea, 2010; Colon et al., 2018). There are, however, some studies that have reported greater contextual fear in females (Fenton et al., 2016; Blume et al., 2017), and several others that have reported no effect or effects that are specific to certain conditions (Pryce et al., 1999; Keiser et al., 2016; Parsons and Russo, 2021; Trott et al., 2022).

These discrepant findings underscore the need for more work to understand the variables that dictate whether sex differences in fear conditioning are observed.

The results from our analyses of the data disaggregated by sex largely confirm the findings from the overall analysis. In both males and females, context conditioning with delayed shock resulted in higher levels of freezing during the context test compared to rats trained with immediate shock. When tested to the light CS used in visual fear conditioning there were no differences across the different intervals or in the type of training received during contextual fear conditioning. However, there was a significant interaction between training type and interval in female rats, suggesting the presence of a time-dependent effect of contextual fear conditioning on the facilitation of subsequent learning. In males, there was no interaction between factors. We also computed correlations between levels of freezing during the context test and levels of FPS to the light during testing. We were interested in understanding the relationship between levels of fear to the two cues as our earlier findings (Lee et al., 2018) showed an inverse relationship between levels of fear to an auditory cue and light cue when rats had received auditory fear conditioning 24 hours before visual fear conditioning. Like this finding, we observed a significant negative correlation in male rats that were trained with a 24-hour interval between the two training episodes. While on average we often see facilitation of learning at this interval (Parsons and Davis, 2012; Parsons et al., 2016; Lee et al., 2018; Cole and Parsons, 2023), these data lend additional support to the idea that there is an interplay between the strength of initial learning and the subsequent effect on new fear learning at 24 hours.

In conclusion, this study examined whether contextual fear conditioning, in both male and female rats, would facilitate learning when rats later encounter a single pairing of light and shock. Our main finding is that contextual fear conditioning does not support the facilitation of later cued visual fear conditioning at any time point, as there were no differences between rats trained with delayed shock, which supports contextual fear learning, and immediate shock which does not support contextual fear. Females exhibited lower levels of cued and contextual fear, indicating either a general deficit in fear learning or lower levels of fear expression in females. These findings provide additional knowledge about the factors that support the ability of prior fear conditioning to facilitate subsequent fear learning and about differences between sexes.

